# GWIS: Genome Wide Inferred Statistics for non-linear functions of multiple phenotypes

**DOI:** 10.1101/035329

**Authors:** H. A. Nieuwboer, R. Pool, C. V. Dolan, D. I. Boomsma, M. G. Nivard

## Abstract

Here we present a method of genome wide inferred study (GWIS) that provides an approximation of genome wide association study (GWAS) summary statistics for a variable that is a function of phenotypes for which GWAS summary statistics, phenotypic means and covariances are available. GWIS can be performed regardless of sample overlap between the GWAS of the phenotypes on which the function depends. As GWIS provides association estimates and their standard errors for each SNP, GWIS can form the basis for polygenic risk scoring, LD score regression^1^, Mendelian randomization studies, biological annotation and other analyses. Here, we replicate a body mass index (BMI) GWAS using GWIS based on a height GWAS and a weight GWAS. We proceed to use a GWIS to further our understanding of the genetic architecture of schizophrenia and bipolar disorder.

---

An example of a well known variable that is a (non-linear) function of multiple phenotypes is BMI, which is defined as weight over height squared. We demonstrate the accuracy of GWIS by reconstructing a body mass index (BMI) GWAS based on publicly available height and weight GWAS summary statistics^2^ (see **URLs**). For each single nucleotide polymorphism (SNP) included in the height and weight GWAS with a minor allele frequency (MAF) larger than 0.05 (as obtained from the HAPMAP Consortium^3^, see **URLs**), we infer estimates and standard errors of these estimates for the association between the SNP and BMI. In a GWAS, BMI must be ascertained for all participants, whereas in a GWIS, we rely on population parameters which reflect the genetic effects on height and weight. Furthermore, the original GWAS for height and weight do not have to be performed in a common set of individuals.

Based on the summary statistics of GWAS for standardized male height and weight^2^, our GWIS replicated 310 out of 356 genome wide hits (an 87.1% replication rate), and found three false positive results (see **Supplementary Table 1**), when compared to a true BMI GWAS performed in the same sample. To demonstrate the method when the constituent phenotypes (i.e., weight and height) are measured independently, we substituted the male height GWAS results for the female height results. Here we assumed the male and female genetic architecture for height in males and females are identical^4^. The GWIS based on independent samples replicated 135 out of 356 genome wide significant signals (a 37.9% replication rate) and yielded eight false positive associations (see **Supplementary Table 2**). All false positives that arise in the female GWIS occured for SNPs which were measured in a small subset of participants (*N* = 1666, where the total sample included up to 73137 women). The Manhattan plots in **Figure 1** revealed that even though there is a loss of power, both forms of GWIS and the original BMI GWAS implicate associations in the same genomic regions.

**Figure 1:**
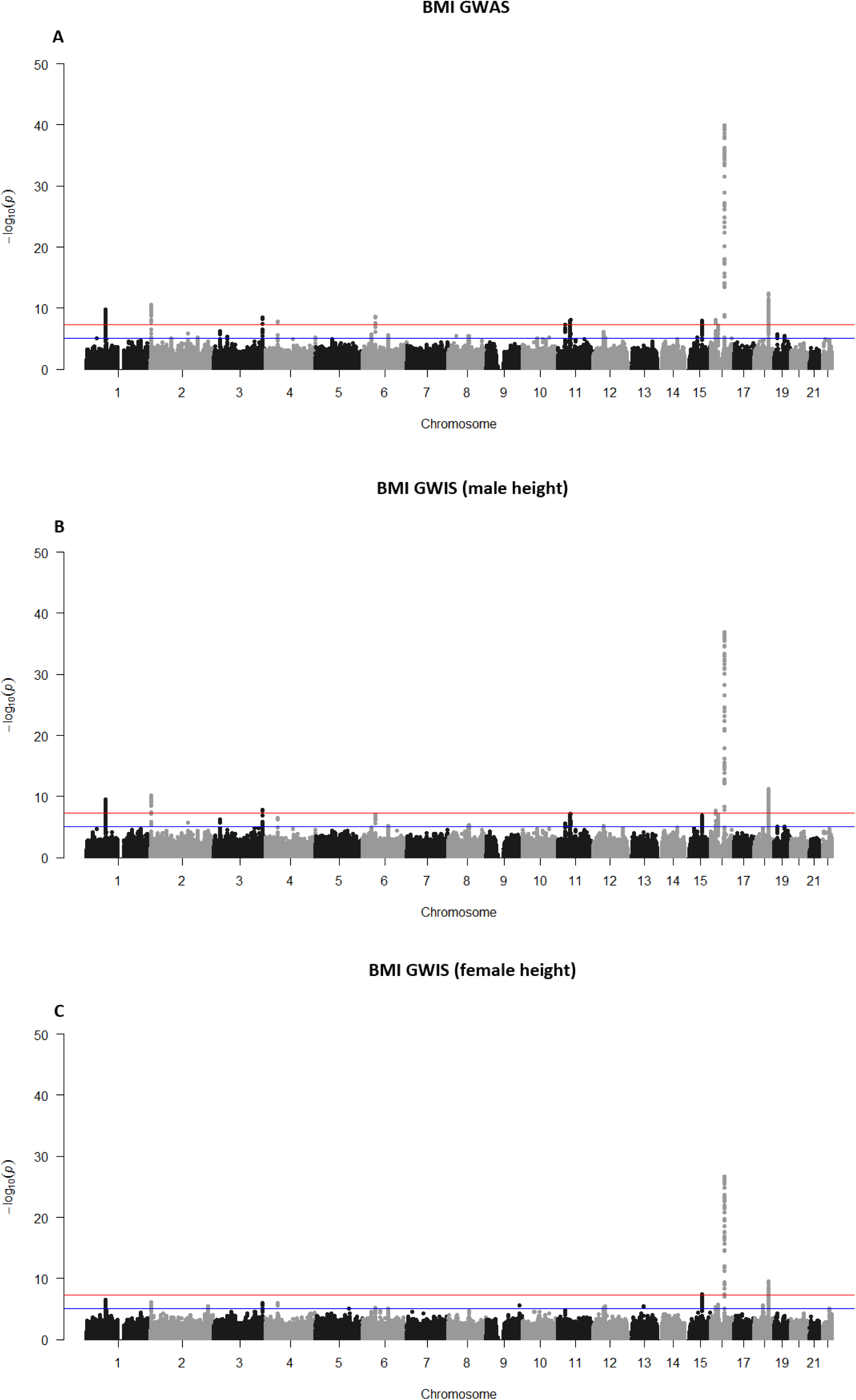
A is a Manhattan plot of − log *p*-values for the BMI GWAS as performed by Randall et al., B is a Manhattan plot of − log *p*-values for the BMI GWIS using male height data, whereas C is a Manhattan plot of − log *p*-values for the BMI GWIS using female height data. The location on the x-axis corresponds to the genomic location of the SNP. In each figure, the blue line corresponds to *p* = 1 · 10^−5^, whereas the red line corresponds to *p* = 5 · 10^−8^.

Using LD score regression^1^, we computed the genetic correlations between BMI based on the GWAS summary statistics, the GWIS using male height data and the GWIS using female height data. As LD score regression requires information on the number of participants available per SNP, we assume the sample size for the BMI GWIS to be the lowest per-SNP sample size of either the height or weight GWAS used. As expected, the genetic correlation between BMI as measured in GWAS, BMI as approximated in GWIS using male height data and BMI as approximated in GWIS using female height data is close to unity (see **Table 1**). Next, we estimated genetic correlations between BMI based on the GWAS, BMI based on GWIS using male height data, BMI based on GWIS using female height data and educational attainment^5^, LDL cholesterol^6^, age at menarche^7^, rheumatoid arthritis^8^ and coronary artery disease^9^. Inference made on the genetic correlates of BMI based on GWIS closely mirror the inference made based on BMI GWAS summary statistics.

**Table 1:** The table reports estimated genetic correlations along with their standard errors between BMI based on GWAS, GWIS using male height data, GWIS using female height data and other traits. These correlations are obtained with LD score regression.

|  | GWIS (female height) | GWAS | Rheumatoid arthritis | Age at menarche | LDL | Educational attainment | Coronary artery disease |
| --- | --- | --- | --- | --- | --- | --- | --- |
| GWIS (male height) | 0.967 (0.012) | 1.007 (0.002) | 0.029 (0.045) | -0.338 (0.035) | 0.019 (0.058) | -0.145 (0.053) | 0.153 (0.063) |
| GWIS (female height) | - | 0.974 (0.013) | 0.018 (0.053) | -0.371 (0.041) | -0.003 (0.061) | -0.157 (0.062) | 0.150 (0.071) |
| GWAS | - | - | 0.039 (0.042) | -0.332 (0.032) | 0.013 (0.050) | -0.160 (0.047) | 0.173 (0.061) |

Ruderfer et al.^10^ performed GWA studies of bipolar disorder (BIP), schizophrenia (SCZ), the pooled bipolar and schizophrenia cases versus the pooled controls (BIP + SCZ) and a GWAS in which the bipolar cases featured as controls and the schizophrenia cases as cases (SCZ − BIP) (see **URLs**). The latter two studies can be reproduced with a GWIS. However, the primary interest of these studies is identifying overlap and contrast between SCZ and BIP. SCZ and BIP are two psychiatric disorders with a substantially correlated genetic underlying liabilities^11^. This correlation prohibits the investigation of genetic variants that are specifically linked to either SCZ or BIP, as well as the investigation of genetic overlap between tertiary traits and SCZ or BIP. As a more exotic application of GWIS, we determine whether the genetic correlation between SCZ or BIP and a tertiary trait is specific to either SCZ or BIP. To this end, we defined a function that decomposes the genetic SCZ liability into a part shared with the genetic liability of BIP and a residual, referred to as unique genetic SCZ liability (unique SCZ). In a similar manner, we defined a function that decomposes the genetic BIP liability into a part shared with the genetic liability of SCZ and a residual, referred to as unique genetic BIP liability (unique BIP). These functions are given by

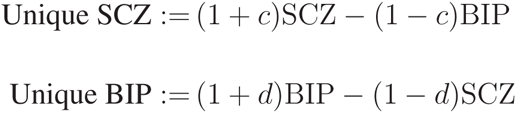

where

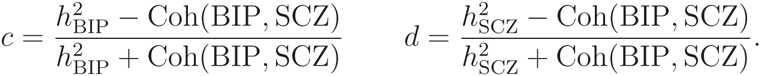

Here, Coh(BIP, SCZ) denotes the coheritability between BIP and SCZ (i.e., *h*_SCZ_ · *r*_BIP,SCZ_ · *h*_BIP_ with *r*_BIP,SCZ_ the latent phenotypical correlation between bipolar disorder and schizophrenia) and 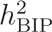, 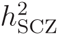 denote the heritabilities of BIP and SCZ, respectively. For the derivation of *c* and *d*, see the online methods. Note that we do not measure unique SCZ or unique BIP in individuals. Furthermore, note that the functions themselves depend on estimated heritability and coheritabilities, which leads to less accurate estimates of genetic effects on unique SCZ and unique BIP. As effect sizes for SCZ and BIP are reported in terms of odds ratios, we take their logarithms to obtain effect sizes on the liabilities.

We performed a GWIS of unique SCZ and a GWIS of unique BIP. For our analysis of unique SCZ and unique BIP in a GWIS, we include SNPs with information values between 0.9 and 1.1 as reported by Ruderfer et al., and minor allele frequencies larger than 0.05 (as obtained from the HAPMAP Consortium^3^), both inclusion criteria reflect common practice in GWA studies^12^. LD score regression^1^ was used to estimate genetic correlations between unique SCZ, unique BIP and educational attainment. We validated the absence of the genetic correlations between unique BIP and SCZ, and unique SCZ and BIP, by applying LD score regression (**Table 2**). Further investigation revealed that unique SCZ does not genetically correlate with educational attainment, whereas unique BIP genetically correlated with educational attainment. This suggests that the observed genetic correlation between schizophrenia liability and educational attainment is fully explained by its genetic correlation with bipolar disorder liability.

**Table 2:** The table reports estimated genetic correlations along with their standard errors between schizophrenia, bipolar disorder, unique schizophrenia, unique bipolar disorder and educational attainment. These correlations are obtained with LD score regression.

|  | Unique SCZ | Unique BIP | SCZ | BIP |
| --- | --- | --- | --- | --- |
| **Educational attainment** | 0.041 (0.082) | 0.218 (0.102) | 0.148 (0.050) | 0.273 (0.067) |
| **BIP** | 0.106 (0.110) | 0.816 (0.031) | 0.572 (0.063) | - |
| **SCZ** | 0.882 (0.026) | -0.016 (0.101) | - | - |

As shown above, GWIS can yield significant novel insight in variables that can be expressed as (non-linear) function of phenotypes. A GWIS can, for example, be performed for equations describing the steady state kinetics of (bio-)chemical reactions involving metabolites of which the concentrations have been analyzed in GWAS, or for equations describing (active) membrane transport of proteins or metabolites given that GWAS summary statistics are available for their concentrations on both sides of the barrier. Other applications of GWIS include converting a GWAS performed for a phenotype as expressed on a logarithmic scale, to a GWIS of the phenotype on its original scale. This transformation can be of use if polygenic risk scoring on the original scale is preferred.

Successful application of GWIS depends on the availability of sufficiently accurate GWAS summary statistics, the number of phenotypes involved in the function, as well as the degree of approximation. The accuracy of the summary statistics of each of the individual GWAS affects the accuracy of the GWIS results. Furthermore, the error of the GWIS statistics increase as more phenotypes are included, due to accumulation of the error in the GWAS results of each of these phenotypes. The degree of approximation used also affects the GWIS results, as the quadratic approximation of a function generally fits better than a linear approximation (see **Supplemental Note 1** for a quadratic approximation of BMI). As the sample sizes used in GWA studies increases, GWIS becomes applicable to a broader domain of functions and yields more accurate results. Related to this last observation, all false positive associations found in the BMI GWIS based on female height data were attributable to a limited sample size for these particular SNPs. We recommend removing SNPs with low allele frequencies, poor imputation quality and SNPs available for a limited number of participants in the original GWA studies before performing GWIS.

With these points of care in mind, however, our method provides a means of obtaining the GWAS summary statistics of a variable that is a function of phenotypes when GWAS summary statistics for these phenotypes are available in (not necessarily overlapping) samples, as outlined in **Figure 2**. This remains possible even when this variable is difficult or impossible to measure in individual participants.

**Figure 2:**
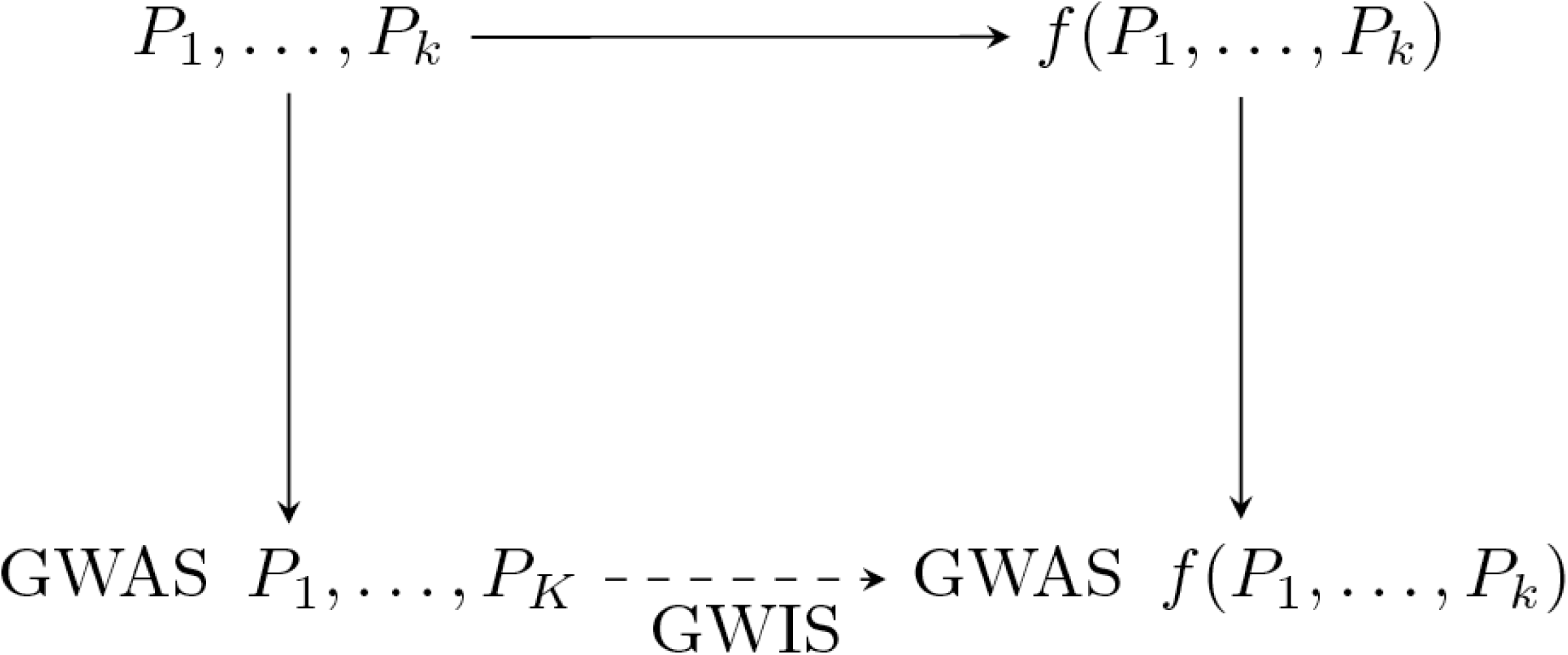
A schematic representation of the role of GWIS in relation to traditional GWAS. GWIS provides a connection between the several GWA studies of phenotypes and a GWA study of a function of these phenotypes, without requiring access to the actual phenotypical data.

## Online methods

Let *V* = *f*(*P*_1_,…*,P_k_*) be a function of the *k* phenotypes *P*_1_,…*,P_k_*. Furthermore, let *S* ~ bin(*n* =2*, p* = effect allele frequency (EAF)) be a binomially distributed variable corresponding to the number of effect alleles (EA) of a biallelic SNP. Let *N* denote the sample size. We assume we have a multivariate linear regression model

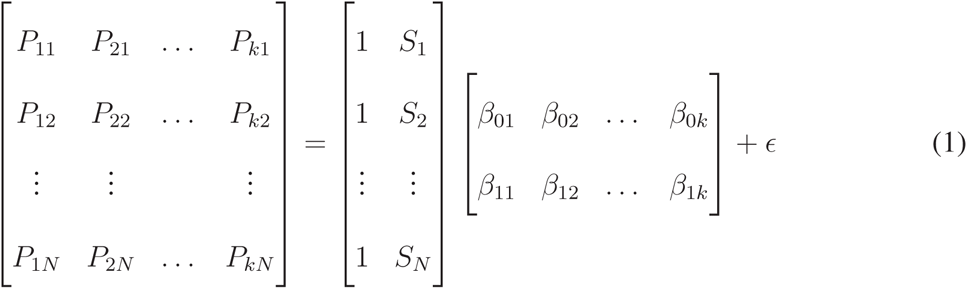

which we write as

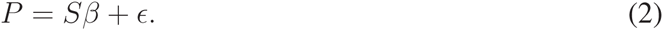

*P* is a *N* × *k* matrix, *S* is a *N* × 2 matrix, *β* is a 2 × *k* matrix and *ɛ* is a *N* × *k* matrix. We assume *ɛ* is a matrix where the columns are normally distributed with zero mean and covariance matrix Σ. Only an estimate for the matrix *β* called 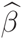 is known, along with the standard errors of each of the 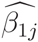, the covariance matrix between the phenotypes *P*_1_,…*,P_k_* and the mean of each phenotype. This is equivalent to having the summary statistics of the GWA studies of each of the *k* phenotypes and their phenotypic covariances.

The goal is to estimate *λ*_0_*,λ*_1_ in

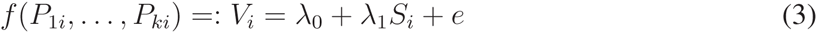

with *e* normally distributed with zero mean. This is equivalent to performing a GWAS of *V*. To do this, we use a first-order Taylor approximation of *V* around the point

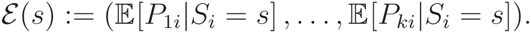

The point 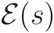 corresponds to the mean of the phenotypes of the individuals that have *s* effect alleles on this SNP. The first-order Taylor approximation is of the form

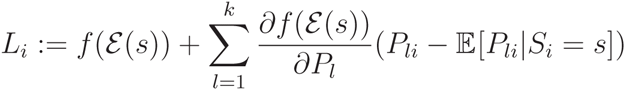

where 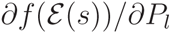 denotes the partial derivative of *f* with respect to *P_l_*, evaluated in the point 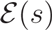. Then, it follows that

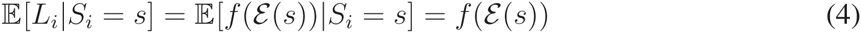

since for each *l* in 1,…*,k*,

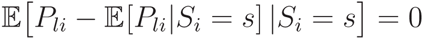

by the linearity of the expectation operator. Equation (4) shows that the mean of the linear approximation is equal to the function evaluated in the phenotypic mean of individuals that have *s* effect alleles. The error incurred in the linearization process takes the form

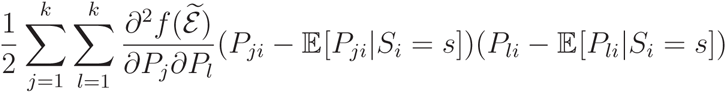

for some 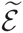 inbetween the two points (*P*_1_*_i_*,…*,P_ki_*) and 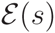.

Note that the linearization is only possible if *f* satisfies certain regularity conditions on the relevant space of phenotype values. For example, division by 0 is not allowed. This can be avoided by linearly transforming the observed phenotypes, along with their associated parts of the *β*-matrix.

We now attempt to derive a linear model for our approximate expression for 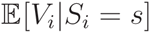. We write

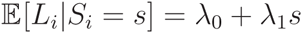

and note that if *s* is 0, we have a direct approximation for *λ*_0_:

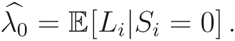

However, as we have shown, 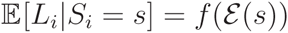, so our approximation for *λ*_0_ becomes

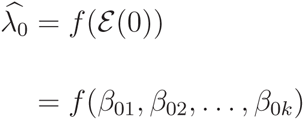

i.e., the function *f* evaluated at the intercepts of our linear regression model. We can also estimate 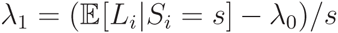 by evaluating this expression for *s* = 1,2 and weighing the results by their (estimated) relative population frequencies. The expression for 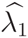 is given by

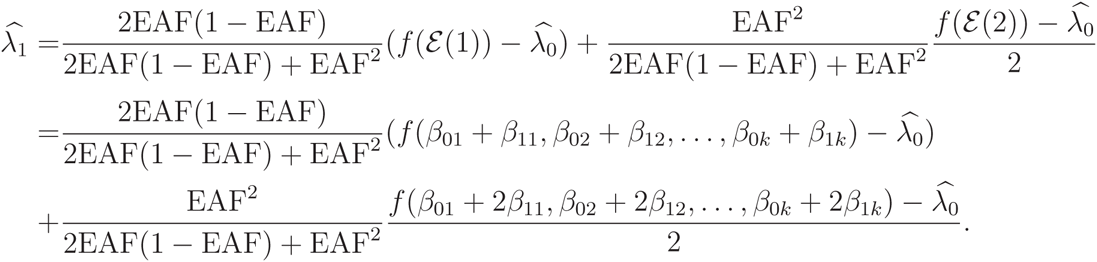

To test our estimates for *λ*_0_ and *λ*_1_, their standard errors must be obtained. However, since we do not have the covariance matrix of 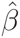, we must first estimate the covariance between each of the 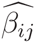. With the theory of multivariate linear regression, we know that the least squares solution to the model *P* = *Sβ*+ *ɛ* is given by

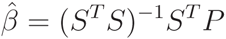

with corresponding variance-covariance matrix

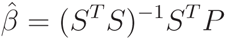

assuming that columns of *ɛ* have zero mean and the rows of *ɛ* are pairwise uncorrelated^13^. This is under the assumption of complete sample overlap. The matrix Σ is a *k* × *k* matrix with the elements Σ*_jl_* =Cov(*ɛ_j_*, *ɛ_l_*), the covariance between the errors in the linear regressions of the phenotypes *P_j_* and *P_l_* on *S*. We assume that the effect of each of the individual SNPs is small, so Var(*ɛ_j_*) ≈ Var(*P_j_*) and Cov(*ɛ_j_*, *ɛ_l_*) ≈ Cov(*P_j_*, *P_l_*). Expanding *S^T^ S* gives

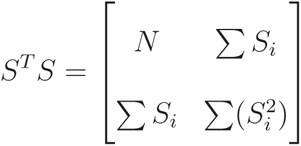

with inverse

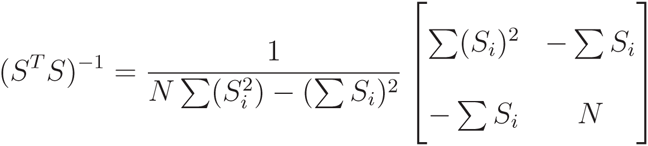

From this, we can infer

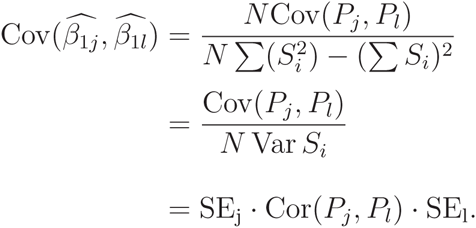

In case there is only partial sample overlap, 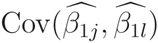 may also be approximated as

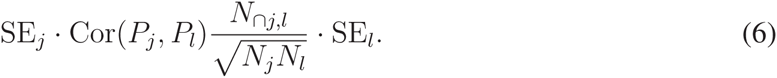

Here, *N*_∩*j*,*l*_ is the number of individuals that is present in both the GWAS of *P_j_* and the GWAS of *P_l_*, *N_j_* is the number of individuals in the GWAS for *P_j_* and *N_l_* is the number of individuals in the GWAS for *P_l_*. If one cannot determine Cor(*P_j_*, *P_l_*) directly or the sample overlap between the GWA studies is unknown, it is possible to use LD score regression based on the summary statistics to estimate 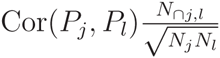. Note that in the absence of sample overlap, *N*_∩*j*,*l*_ is zero and thus 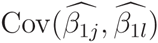 is zero.

Having obtained the covariance matrix for 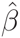, we can apply the Delta-method^14^ to find the standard errors of 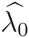 and 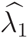. The derivation above is done in terms of linear regression assuming a continous response variable. However, a link function may be used to apply this to other response variables.

Here, we outline a GWIS as applied to BMI. BMI is defined as weight (in kilograms) divided by height (in meters) squared. Let *μ_w_*, *μ_h_* denote the means of respectively weight and height and let *α_w_*, *α_h_*, *β_w_*, *β_h_* denote the intercepts of weight and height and the regression coefficients in the regression of weight and height on the SNP respectively. We assume all of these parameters are known. As shown above, the mean of our approximated BMI is equal to

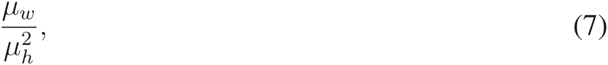

i.e., BMI calculated for the mean weight and mean height. In our case, the GWA summary statistics were for standardized weight and height, but were destandardized before computing the GWIS. The destandardization is based on information on population averages and standard deviations obtained from the Netherlands Twin Register (NTR)^15^. The destandardization involves multiplying the effect sizes by the standard deviation and using the population mean as a substitute for the intercept. The mean of the appromixation is in general going to be equal to the function evaluated in the means of the phenotypes. The linear regression of BMI on the number of effect alleles of a given SNP is

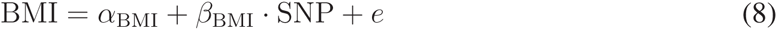

where *α*_BMI_ is the intercept of the linear regression, *β*_BMI_ is the regression coefficient and *e* is the error of the linear regression.

Then, the derived values for the intercept and the regression coefficient become

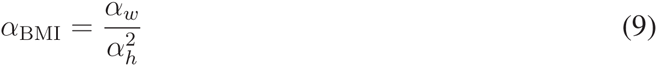

and

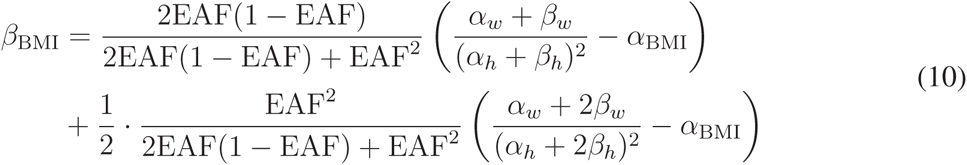

where EAF is the effect allele frequency of the SNP.

In our examples, we have used a linear approximation to perform the GWIS; however, in **Supplemental Note 1** we outline the second order approximation of BMI, which should be used in conjunction with a second order Delta-rule.

Given two phenotypes *A* and *B*, we can use our method to define a new trait as

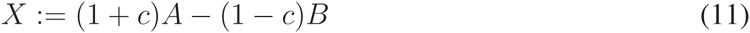

for a specific constant *c*. This constant is chosen such that a certain type of correlation between *X* and *B* becomes zero and the correlation between *A* and *X* is nonzero. Note that this correlation may be genetic, environmental or phenotypic, depending on the application. In terms of linear regression, this can be seen as

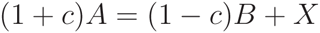

so that *X* is the residual of the linear regression (with fixed coefficients) of (1 + *c*)*A* on (1 − *c*)*B*.

Note that zero correlation does not imply that *X* and *B* are independent; rather, they have only become linearly independent. The expression for *c* is

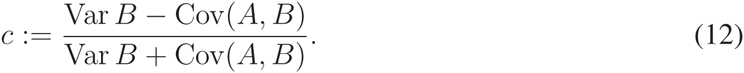

Note that Cov and Var here denote the covariances and variances that are specific to the type of correlation that is considered. For example, in the case of genetic correlation, Cov denotes the coheritability and Var denotes the heritability of the traits.

We derive *c* by solving Cov(*X*, *B*) = 0, so

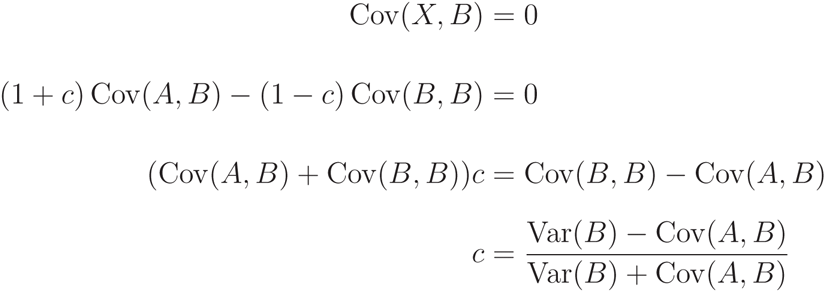

which is well-defined if and only if Var(*B*) ≠ − Cov(*A*, *B*), that is, *B* is not equal to −*A*.

An equivalent expression for *c* is

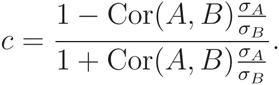

The term 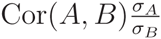 corresponds to the slope of the linear regression of *B* on *A*. Thus, *X* is actually the distance between the data points in a 2-dimensional plane and their projection onto the linear regression line of *A* on *B*, rather than the vertical distance between the predicted value of *A* and the data point. This allows for error in the assessment of both *A* and *B*, rather than only measurement error in *A*. This is important since *X* is analyzed in a GWIS and the estimates for the association between both *A* and *B* and a SNP have a certain standard error.

## Supplemental Note 1. A second order approximation of BMI

The second order Taylor approximation of BMI is given by

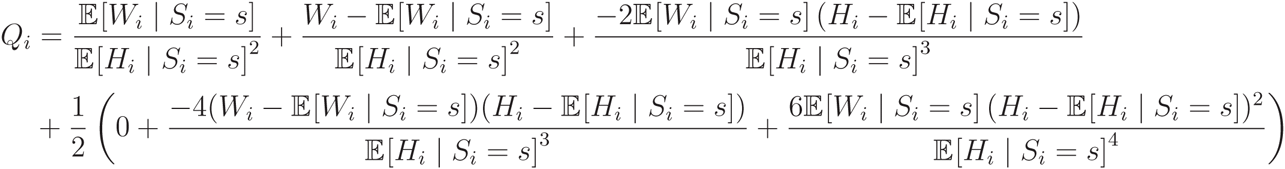

so that

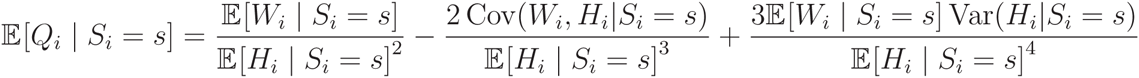

Then, using this for our linear regression

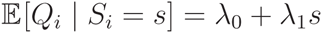

together with *s* = 0 gives

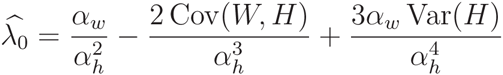

under the assumption that Cov(*W_i_*,*H_i_*|*S_i_* = *s*) = Cov(*W*,*H*) and a similar assumption for the variance of height. Then,

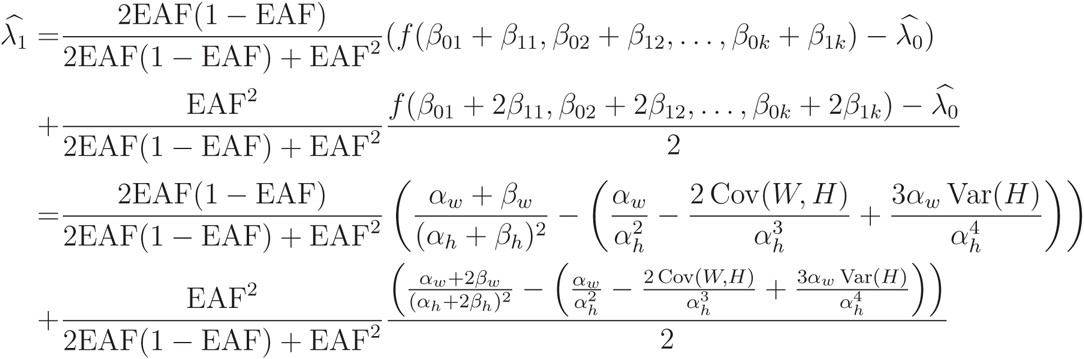

**Supplementary Table 1:**
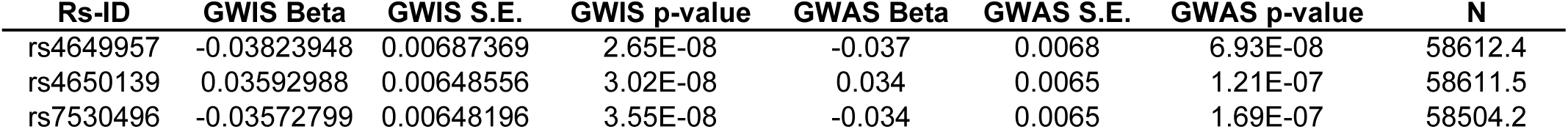
The table reports associations found in the BMI GWIS based on male height data which are not significant in the original BMI GWAS. Note the differences in effect sizes are minor and *p*-values are close to the significance threshold in both analyses.

| Rs-ID | GWIS Beta | GWIS S.E. | GWIS p-value | GWAS Beta | GWAS S.E. | GWAS p-value | N |
| --- | --- | --- | --- | --- | --- | --- | --- |
| rs4649957 | -0.03823948 | 0.00687369 | 2.65E-08 | -0.037 | 0.0068 | 6.93E-08 | 58612.4 |
| rs4650139 | 0.03592988 | 0.00648556 | 3.02E-08 | 0.034 | 0.0065 | 1.21E-07 | 58611.5 |
| rs7530496 | -0.03572799 | 0.00648196 | 3.55E-08 | -0.034 | 0.0065 | 1.69E-07 | 58504.2 |

**Supplementary Table 2:** The table reports associations found in the BMI GWIS based on female height data which are not significant in the original BMI GWAS. Note the strong deviance in effect size, standard error and *p*-value between the analyses. This deviance is likely caused by the limited number of individuals for which these SNPs were measured in the female height GWAS.

| Rs-ID | GWIS Beta | GWIS S.E. | GWIS p-value | GWAS Beta | GWAS S.E. | GWAS p-value | N |
| --- | --- | --- | --- | --- | --- | --- | --- |
| rs11206949 | -5.71393947 | 0.010152767 | 0 | 0.94 | 1.1 | 0.38 | 1666 |
| rs11206950 | -2.91013533 | 0.00155349 | 0 | -0.94 | 1.1 | 0.38 | 1666 |
| rs11206951 | -5.71393947 | 0.010152767 | 0 | 0.94 | 1.1 | 0.38 | 1666 |
| rs11206952 | -5.71393947 | 0.010152767 | 0 | 0.94 | 1.1 | 0.38 | 1666 |
| rs12073104 | -5.71969538 | 0.010170506 | 0 | 0.94 | 1.1 | 0.38 | 1666 |
| rs12080262 | -2.91013533 | 0.00155349 | 0 | -0.94 | 1.1 | 0.38 | 1666 |
| rs12083335 | -2.9083309 | 0.005892014 | 0 | -0.94 | 1.1 | 0.38 | 1666 |
| rs12087759 | -2.91013533 | 0.00155349 | 0 | -0.94 | 1.1 | 0.38 | 1666 |

## URLs

PGC summary statistics used in the schizophrenia and bipolar disorder analysis^10^:

https://www.med.unc.edu/pgc/results

GIANT summary statistics used in the BMI analyses^2^:

https://www.broadinstitute.org/collaboration/giant/index.php/GIANT_consortium_data_files

HAPMAP 2 allele frequencies were obtained from the the public webpage of the HAPMAP Consortium^3^:

http://hapmap.ncbi.nlm.nih.gov/downloads/index.html.en

